# Spontaneous transfer of small peripheral peptides between supported lipid bilayer and giant unilamellar vesicles

**DOI:** 10.1101/2023.06.08.544237

**Authors:** Emanuela Efodili, Ashlynn Knight, Maryem Mirza, Cedric Briones, Il-Hyung Lee

## Abstract

Vesicular trafficking facilitates material transport between membrane-bound organelles. Membrane protein cargos are trafficked for relocation, recycling, and degradation during various physiological processes. *In vitro* fusion studies utilized synthetic lipid membranes to study the molecular mechanisms of vesicular trafficking and to develop synthetic materials mimicking the biological membrane trafficking. Various fusogenic conditions which can induce vesicular fusion have been used to establish synthetic systems that can mimic biological systems. Despite these efforts, the mechanisms underlying vesicular trafficking of membrane proteins remain limited and robust *in vitro* methods that can construct synthetic trafficking systems for membrane proteins between large membranes (>1 μm2) are unavailable. Here, we provide data to show the spontaneous transfer of small membrane-bound peptides (∼4 kD) between a supported lipid bilayer (SLB) and giant unilamellar vesicles (GUVs). We found that the contact between the SLB and GUVs led to the occasional but notable transfer of membrane-bound peptides in a physiological saline buffer condition (pH 7.4, 150 mM NaCl). Quantitative and dynamic time-lapse analyses suggested that the observed exchange occurred through the formation of hemi-fusion stalks between the SLB and GUVs. Larger protein cargos with a size of ∼77 kD could not be transferred between the SLB and GUVs, suggesting that the larger-sized cargos limited diffusion across the hemi-fusion stalk, which was predicted to have a highly curved structure. Our system serves as an example synthetic platform that enables the investigation of small-peptide trafficking between synthetic membranes and reveals hemi-fused lipid bridge formation as a mechanism of peptide transfer.

**Graphical abstracts:** 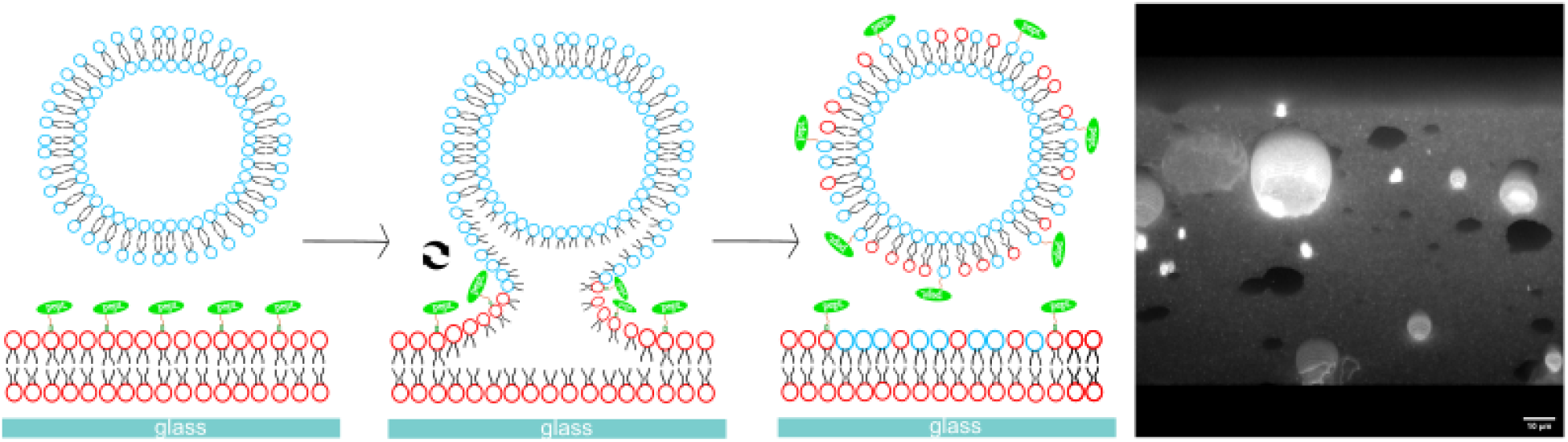

## 1. Introduction

Vesicular trafficking is a process of material transport between two lipid bilayers. In living cells, this occurs between different membrane-bound organelles such as plasma membranes, lysosomes, and endoplasmic reticulum [1-3]. The process often involves vesicular fusion which is an event where two lipid bilayers contact, hemi-fuse, and completely fuse to become a continuous bilayer [4, 5]. Vesicular trafficking is one of the most fundamental activities in living cells, thus malfunction of the trafficking is known to cause various diseases [6, 7]. Naturally, understanding the molecular mechanism of vesicular trafficking has been an intense area of research. Vesicular trafficking by synthetic lipid vesicle systems has been used widely to study the molecular mechanism of the trafficking process in controlled environments [8-10] and also to find applications as biomimetic materials in creation of artificial life [11, 12] and drug delivery systems [13, 14]. Structural and biophysical studies on trafficking systems made significant progress in understanding the molecular mechanism in the last few decades, but we currently still do not have enough understanding and techniques to synthetically reproduce and construct a system capable of performing vesicular trafficking of membrane proteins.

Physiological trafficking may involve several tens of proteins, and each trafficking pathway between different organelles involve unique set of proteins, thus making study of molecular mechanism challenging [1, 15]. Cellular imaging by fluorescence microscopy is often performed to study the localization of different proteins at the sites of trafficking [16-18] and hypothetical mechanism can be tested with purified proteins *in vitro* by reconstitution study [8, 10, 19]. Such studies often focus on the structure-function relationship of the proteins with known hypothetical roles in the process. The studies are crucial for potential drug development targeting the trafficking processes, but physiological trafficking process tends to be very complex, so establishing a simple physical model toward the development of simple synthetic system is not trivial from such studies.

*In vitro* studies to develop synthetic fusion system has been hinting on minimal necessary components for successful vesicular fusion. Such studies often use chemical methods that might not even be physiologically relevant looking for applications of vesicular trafficking as biomimetic materials. Various fusogenic conditions are known to catalyze the fusion or formation of hemi-fusion stalks between lipid bilayers. Example fusogenic conditions include multivalent cations [20, 21], freeze-thawing [22, 23], nano-particle interaction [24, 25], and functional peptide mediated fusions [26, 27] to name a few. Previous reports show examples of vesicular content inside may be transferred to another vesicle by fusion [28, 29]. However, examples on membrane proteins, proteins spanning or peripherally anchored to the lipid membranes, being transferred [26, 30] have been underexplored, limiting our understanding on membrane protein trafficking processes.

In this study, we report a case of synthetic vesicular trafficking of peripheral peptides between supported lipid bilayer (SLB, a lipid bilayer formed on glass surface), and giant unilamellar vesicles (GUV, single layered lipid vesicle with dimeter greater than 1 μm). We used confocal laser scanning microscopy (CLSM) to monitor the exchange of small peptides anchored on the lipid membranes from SLB to GUV. Our results show that such transfer may occur spontaneously between SLB and GUV under the neutral saline condition of pH 7.4 and 150 mM NaCl by hemi-fusion stalk formation between SLB and GUV.

## 2. Methods

### 2.1. GUV Preparation

The GUVs were prepared by gentle hydration [31-33]. This involved combining a lipid mixture of calculated composition in a clean round-bottomed flask using glass syringes (Hamilton Company). The composition of the GUV used in this study was 30% 1,2-dioleoyl-sn-glycero-3-phosphoethanolamine (DOPE), 5% 1,2-dioleoyl-sn-glycero-3-phospho-L-serine (DOPS), 65% 1,2-dioleoyl-sn-glycero-3-phosphocholine (DOPC) by mol% unless specified otherwise. DOPC is a commonly used model lipid for its low melting temperature and fluidity, DOPS is a net negatively charged lipid at pH 7.4 introduced to facilitate vesicle swelling process, DOPE is a lipid that can facilitate flexible deformation of the lipid membranes by curved structure formation without affecting the membrane fluidity [34]. Negative control sample composition was 5% Ni-DGS, 95% DOPC. 0.1% of the DiD (Invitrogen) replaced the DOPC of each composition when fluorescent reporter was used. The mixture was then dried to a thin film with nitrogen gas, to evaporate the chloroform solvent, while on a hot plate set to maintain the temperature at 55-60°C. The flask was placed in a room-temperature vacuum chamber for the sample to dry further for at least one hour. 1 mL of 320 mM sucrose solution was introduced into the flask, the flask was sealed, and incubated overnight at 37 °C to induce vesicle swelling by hydration.

Following incubation, the now GUVs were centrifuged for 5 min at 12,000 g to remove large lipid aggregates. 600-800 μL of aggregate free solution were collected and stored at 4°C until use.

The lipids used were obtained from Avanti Polar Lipids Inc as. High purity laboratory water that was properly purified and deionized was used for the experiments.

### 2.2. SLB Preparation and functionalization

The small unilamellar vesicles (SUVs) were prepared by extrusion [8]. This involved combining a lipid mixture of calculated composition in a clean round-bottomed flask using glass pipettes (Hamilton). The composition of the SUV used in this study was 30% DOPE, 5% Ni bound 1,2-dioleoyl-sn-glycero-3-[(N-(5-amino-1-carboxypentyl)iminodiacetic acid)succinyl] (Ni-DGS), 65% DOPC unless specified otherwise. 0.1% of the Texas Red 1,2-Dihexadecanoyl-sn-Glycero-3-Phosphoethanolamine (TR-DHPE, Invitrogen) replaced the DOPC when fluorescent reporter was used. Ni-DGS serves the role of anchoring peptides and proteins to the membranes by poly histidine-Ni chelation. The mixture was proceeded to dry with nitrogen gas, to evaporate the chloroform solvent, while on a hot plate set to 55-60 °C, to ensure temperature control. The flask was placed in a room-temperature vacuum chamber for the sample to dry further at least for one hour. 1 mL of deionized water was introduced into the flask, covered, then freeze-thawed at least three times using Lab Armor metal beads (Gibco) stored at -80 °C. SUV extrusion was performed using handheld mini extruder nine times using a 100 nm pore-size polycarbonate filter (Whatman). Final SUVs were stored at 4 °C until use. The lipids and handheld mini extruder were obtained from Avanti Polar Lipids Inc.

The SLB was prepared by SUV rupturing. Assembled was an Attofluor chamber (Invitrogen) with a round cover glass (25CIR-1, ThermoFisher), that was sonicated for 30 minutes in solution of isopropyl alcohol:water = 1:1 by volume, then etched with acid piranha solution (30% hydrogen peroxide:concentrated sulfuric acid = 1:3 by volume). A silicon O-ring was placed for a reduced sample space. 100 uL of SUV solution and 100 uL of phosphate buffer including magnesium chloride (20 mM phosphate, 5 mM MgCl_2_, pH 7.4) were combined in an Eppendorf tube. 200 uL of the mixture was dispensed into the chamber and left to incubate for 30 min at room temperature. The SUVs spontaneously ruptured on the piranha-etched glass to generate the SLB [35]. Without exposing the SLB to the air to avoid oxidation of the bilayer, the chamber was rinsed ten times with Hepes buffer (20 mM Hepes, 150 mM NaCl, pH 7.4). When incubating the peptide or protein on the SLB surface, a diluted amount of the peptide/protein concentration (10 nM – 1 μM) was introduced into the chamber and left to sit for 30 min. Linker for activation of T-cell (LAT) peptide [33] and Small Ubiquitin-like modifier 3– Green fluorescence protein (SUMO3-GFP) [31, 36] included his-tags to covalently bind to the Ni of the Ni-DGS lipids. LAT peptide was covalently labeled (Cysteine-maleimide chemistry) with Oregon Green 488 (OG488, Invitrogen) or Alexa Fluor 647 (A647, Invitrogen), and SUMO3-GFP is inherently fluorescent by the GFP fluorescence. Without exposing the SLB to the air, the chamber was then rinsed ten times with Hepes buffer (20 mM Hepes, 150 mM NaCl, pH 7.4).

To ensure stable protein binding, incubation proceeded for another 20 minutes [37], then finally rinsed 7x without exposing the SLB to the air. Fluorescence bleaching after photobleaching (FRAP) measurement was performed to ensure fluidity of the SLB formed (Supplementary S1) [8].

### 2.3. Imaging sample preparation

After checking the creation of fluidic SLB, depending on the experiment and composition, 1-10 uL of GUV solution was added to the chamber with Hepes buffer to ensure an adequate amount of vesicles were observable per image, then settled for 5 min on the microscope stage. This step was where spontaneous interaction between the SLB-peptide (protein) and GUV was observed. Time-lapse movies and z-stack images with multi-color excitation were taken accordingly to observe exchange of lipids and peptides (proteins). At least 30 min of interaction between SLB and GUVs was allowed at room temperature (22°C) before taking the final z-stack images. For the cationic shock experiment, after the initial incubation was complete, an equal volume of 2 mM LaCl_3_ solution in Hepes buffer was gently added to introduce 1 mM La3+ concentration final. The LaCl_3_ triggered the facilitated lipid exchange between the SLB and GUV. Time-lapse movies and z-stack images were taken accordingly. At least 30 min of interaction was allowed before taking the final z-stack images. The sample samples was covered with Petri dish at all times during imaging to prevent sample evaporation.

For the negative control sample, which was to calculate the noise level of co-localization, sample chamber was assembled the same way without doing the piranha etching of the cover glasses. 5 mg/mL BSA (Sigma-Aldrich) solution was used to block the surface for 30 min, followed by addition of GUV solution for image acquisition using the same conditions as the imaging of other samples.

### 2.4. Fluorescence Imaging Conditions

#### 2.4.1. Confocal laser scanning microscopy imaging

Images shown in this report were collected using CLSM unless specified otherwise. Briefly, a Nikon Ti-E-based C2 confocal microscope was used (Nikon, Japan). Excitation laser lights of 488, 561, and 640 nm were used with matching emission filters to collect signals from the fluorescent molecules. A Nikon Plan Apo 100×NA 1.45 oil immersion objective was used without further magnifying the lens in the optical path. The typical mode of scanning was to collect data as 1,024 × 1,024 pixels spanning a 127.3 μm × 127.3 μm area, whereas motorized z-axis movement allowed the automated acquisition of z-stack images. All the example images shown were originally collected by taking a z-stack every 1 μm apart which is typically a great sampling in the z-direction considering the resolution of confocal laser scanning and the size of the GUVs. Most representative image sections showing clear contour of the vesicle fluorescence were chosen for further analysis typically images ∼3-5 μm above the bottom.

#### 2.4.2.. Wide field fluorescence microscopy imaging

A Nikon Ti2E-based inverted epifluorescence microscope system (Nikon, Japan) was used for FRAP and time-lapse imaging. A light-emitting diode (LED) white light excitation source (Lumencor) was optically filtered with multiple optical filters and dichroic mirrors to excite and collect fluorescence emission optimized for Cy5, GFP, and Texas Red, respectively. The Nikon Apo 100X TIRF oil objective with a numerical aperture of 1.49 was used for imaging. Single-molecule sensitivity with a high quantum yields scientific complementary metal–oxide– semiconductor camera was used for data collection (Hamamatsu ORCA Flash 4.0, Hamamatsu, Japan). Automatic z-position control was used for precise z-stack acquisition and x, and y positions were controlled manually. Time lapse images were taken by choosing a z-section(s) of interest to repeat image acquisition at set time intervals (15 s – 60 s). Due to faster acquisition time, wide field fluorescence imaging was often advantageous when observation of dynamic change was the goal. Micromanager (https://micro-manager.org/), an ImageJ (https://imagej.nih.gov/ij/) based software was used to manage the devices. All data collection was performed on a vibration isolation table.

### 2.6. Peptide and protein purification

LAT peptide, a derivative from the physiological LAT was synthesized as a service of the company Biomatic and labeled with OG488 and A647 each appropriately following the manufacturer’s recommended protocols using Cys-maleimide chemistry [33]. The peptide was chosen as a small model peptide in this study, not due to its specific sequence. The sequence of the LAT peptide was:

HHHHHHHHHHGGSGGSGGD[pY]VNVGGSGGSGGD[pY]VNVDKKC

SUMO3-GFP was purified by *E. coli* overexpression following the suggested overexpression condition from the original manuscript that it was developed [36]. A series of chromatography in a fast protein liquid chromatography system (GE Healthcare) was performed [31]. The plasmid was given by the Michael Rosen group (Addgene plasmid #127093).

### 2.7. Data analysis

Images were analyzed using a home-built automated script. (Run in the freely available software GNU Octave 6.4.1, https://octave.org/, full code available in the Supplementary S4) In our experiments, GUV lipids, SLB lipids, and peripheral peptides (proteins) were labeled by fluorescent dyes with different wavelengths of excitation/emission. We used binary masking and pixel counting to quantify the degree of pixel colocalization between different channels.

Colocalized pixels mean coexistence within the same lipid membrane thus serves the purpose of quantifying the degree of lipids/peptides exchange. By doing this, potential ambiguity of using raw intensity values, which may vary at different optical setups, could be minimized. One of the z-stack images, typically ones 3-4 μm above the SLB surface was chosen to perform binary masking in a way that typical level of noise signal was excluded (count as 0) and net signal from the membranes were counted (count as 1). Number of overlapping or colocalizing pixels in multiple channels were counted for quantification. For quantification of peptide (protein) transfer, DiD channel, or the channel representing lipids originally included in the GUV were used as a reference channel and OG488 (GFP) channel, the channel representing the fluorescence of peptides (proteins) were counted for overlapping pixels to report ratio of [Number of pixels colocalized in OG488 channel] / [Number of pixels representing GUV lipids by DiD]. For lipid exchange analysis, the ratio of [Number of pixels colocalized in TR-DHPE channel representing SLB lipids] / [Number of pixels representing GUV lipids by DiD] was reported. To selectively analyze the vesicles that went through the peptide transfer, only pixels colocalized with the OG488 channel was used for quantification.

## 3. Results and discussion

### 3.1. Small membrane bound peptides spontaneously transfer between supported lipid bilayer and giant unilamellar vesicles

Figure 1A shows a schematic of the peptide exchange experiment between SLB and GUV. Our original goal was to create a synthetic system capable of trafficking membrane peptides between SLB and GUV catalyzed by known fusogenic conditions such as cationic shock, glass nanoparticles, and freeze thawing [20-25, 38]. During the investigation, we unexpectedly learned that spontaneous peptide transfer between SLB and GUV occurs reproducibly under the condition of neutral physiological saline buffer (20mM Hepes, 150mM NaCl, pH 7.4) that we decided to study in details.

**Figure 1.**
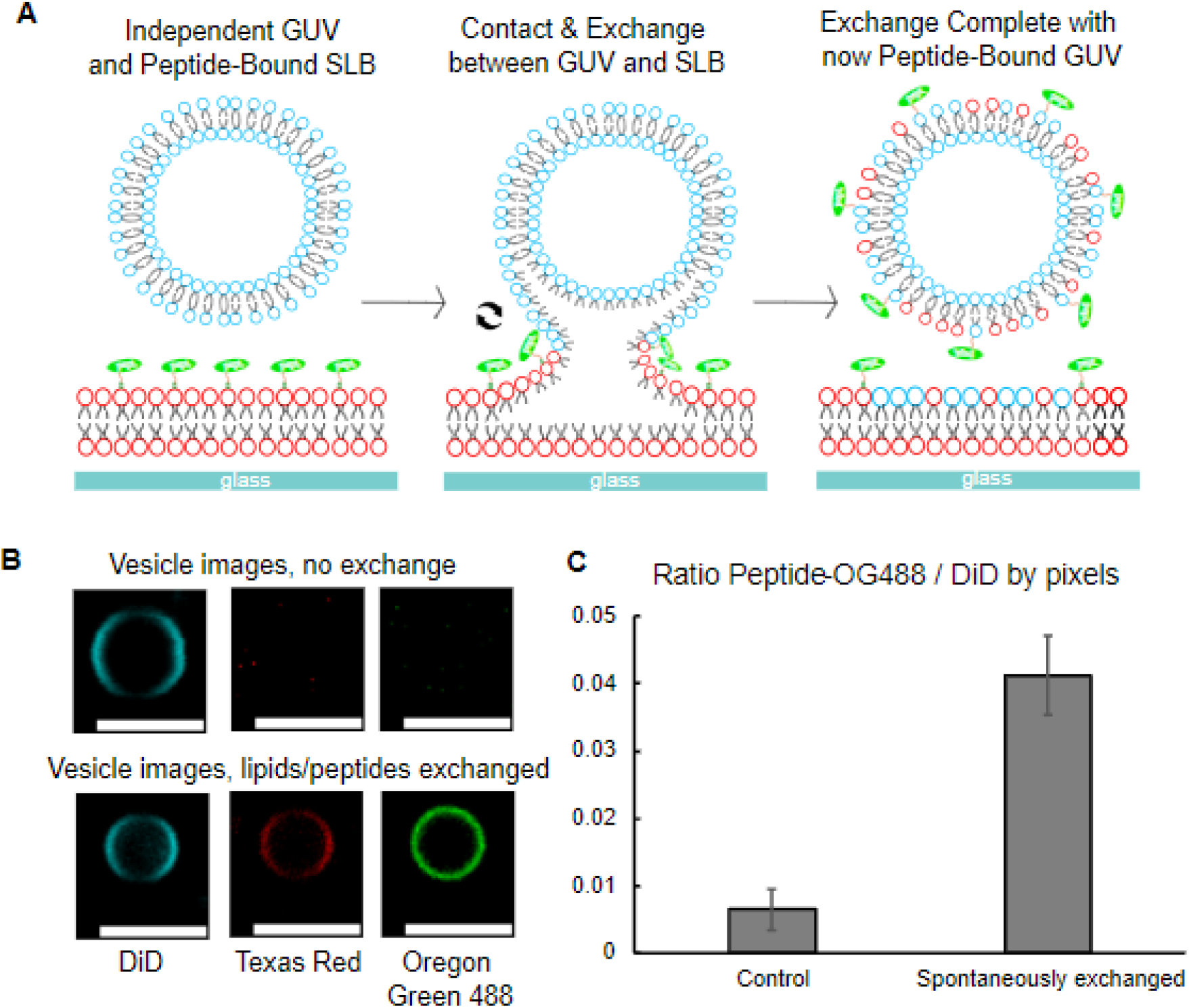
Phospholipid membrane fusion and protein transfer from SLB to GUV. (A) Schematic of the SLB to GUV exchange with peptide anchored onto the SLB. The lipid composition for the GUV was 64.9% DOPC, 30% DOPE, 5% DOPS, and 0.1% DiD. The lipid composition for the SLB was 64.9% DOPC, 30% DOPE, 5% Ni-DGS, and 0.1% TR-DHPE. (B) Example images of DiD, TR-DHPE, and peptide-OG488 fluorescent vesicles when the exchange does and does not occur. Scale bars are set to 5 μm. (C) Statistical distribution by pixels of the ratio of LAT-OG488 to DiD in a negative control setting compared to spontaneous exchange, where spontaneous exchange occurs at an observable rate. Error bars represent standard errors of multiple images from multiple samples.

To test the peptide transfer, SLB was formed on piranha-etched coverglass (30% DOPS, 5% Ni-DGS, 54.9% DOPC, and 0.1% DiD by mol%). Ni-DGS was used to covalently anchor fluorescently labeled small peptides (LAT peptide derived from the mammalian T-cell signaling protein [33, 39]) to the SLB using his-tags. GUVs were introduced by gentle pipetting and the state of peptide exchange was monitored by CLSM images. SLB lipids, GUV lipids, and peptides were each labeled with different wavelengths of fluorescence reporters in order to monitor the change of lipid membrane morphology and exchange of molecules by fluorescence image analysis. Multiple images were collected after 30 min of interaction between SLB and GUV, and images were analyzed for quantification.

Representative images for the case of peptide exchange is shown in Figure 1B. Green fluorescence from LAT-OG488 is clearly observed in the GUV suggesting the transfer of the peptides. We quantified the degree of exchange by calculating the number of colocalizing pixels in the image with peptide fluorescence divided by the total number of pixels with lipid fluorescence. Figure 1C shows the quantitative comparison of peptides on the membranes between the negative control (which only included DiD lipid fluorescence reporter) and the peptide exchanged vesicles. The ratio below 0.01 of the control sample represents the noise level. As shown in the quantification, exchange of peptide is clear although not all lipids membranes ended up taking up the peptides from the SLB. 0.04 of ratio might seem relatively low, but the case of peptide exchange was very clearly observable by fluorescence, evenly distributed in different regions of the samples, and reproducible. It shows that certain ratio of

GUV-SLB contact interactions led to spontaneous exchange of membrane bound peptides under the physiological ionic strength and pH, and not all contact interactions led to the successful exchange of peptides.

### 3.2. Peptide exchange between membranes involve lipid exchange

We hypothesized that membrane peptide transfer occurs as a result of natural lipids exchange between SLB and GUV. The lipid intensities of vesicles that showed peptide transfer were analyzed to monitor the degree of lipid exchange using the pixel number analysis similar to

Figure 1. Number of pixels with TR-DHPE (lipids originally in SLB) over number of pixels with DiD (lipids originally in GUV) was quantified only for pixels with LAT-OG488 as a result of peptide exchange. The quantitation indeed showed that vesicles that took up the peptides did exchange lipids evidenced by 0.70 of the TR-DHPE/DiD ratio (Figure 2A). It means most lipid membranes that took up peptides had lipid fluorescence from reporters that were originally in GUVs and in SLB both. Such process of lipid exchange often involved observable diffusion of reporter fluorescence, originally in GUV, across the SLB surface showing that the process does involve thermal diffusion (Supplementary S2). Also notable is that DiD fluorescence pixel counting usually did not reach zero after the exchange. It means a certain amount of DiD reporters, originally in GUVs still remained after the interaction. This observation supports the idea that the exchange occurs by hemi-fusion involving the fusion of only one leaflet instead of the full fusion of both leaflets which would inevitably lead to depletion of DiD fluorescence in GUV after the lipid exchange. This is because the amount of lipids in each vesicle is limited compared to the amount of lipids in SLB which is a fluidic bilayer spanning the entire cover glass. If only one of the two leaflet was to hemi-fuse, fluorescence reporters in the unfused leaflet of the vesicles would still remain. Overall, the analysis suggests that the peptide transfer involves lipid transfer between two fluidic membranes and the observed peptide exchange is not an event that is only specific to the peptides.

**Figure 2.**
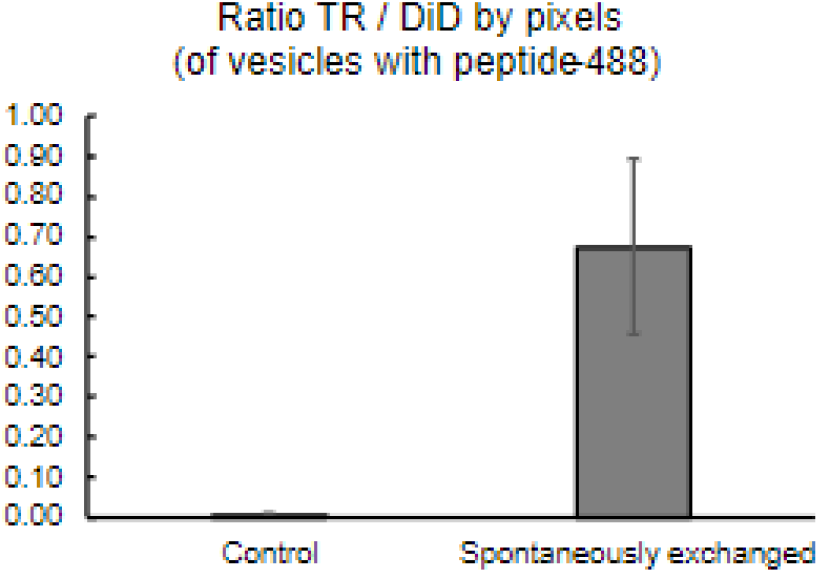
Statistical distribution by pixels of the ratio of TR-DHPE vesicles with LAT-OG488 to DiD vesicles with LAT-OG488. There is an observable increase in TR-DHPE to DiD vesicles that spontaneously exchanged peptides. Error bars represent standard errors of multiple images from multiple samples.

### 3.3. Time lapse images show that the peptides are transferred by occasional hemi-fusion bridge formation mechanism

To better monitor the dynamic process of the peptide and lipid exchange, we used different sets of fluorescent reporters and took time lapse movies at the contact site between the GUV and SLB using wide field fluorescence microscopy. Figure 3 shows example time-lapse movies at the contact site and z-section images after the exchange. Figure 3AB images were taken by fluoresce reporters of TR-DHPE in SLB, A647 on peptides, and Figure 3C images were taken by fluorescence reporters of A647 on peptides only. Figure 3AB show progress of most typical modes of peptide exchange and Figure 3C shows an example images of the sample after the peptide exchange happened. Before GUVs contacted SLB, peptide fluorescence was evenly distributed showing homogeneous distribution of peptides on the SLB. As shown in Figure 3A, once GUVs landed to make contact, dark peptide fluorescence became visible suggesting peptides were being excluded at the contact site. Such exclusion was never observed in the lipid channel suggesting that this was physical exclusion of peptides at the contact site due to proximity of two lipid membranes. This dark spot was followed by intermittent peptide fluorescence among the dark spot, and soon the colorless GUVs took up the fluorescence of both lipids and peptides simultaneously suggesting exchange of lipids and peptides. The GUVs that used to be invisible due to lack of fluorescence reporters became clearly visible at this point.

**Figure 3.**
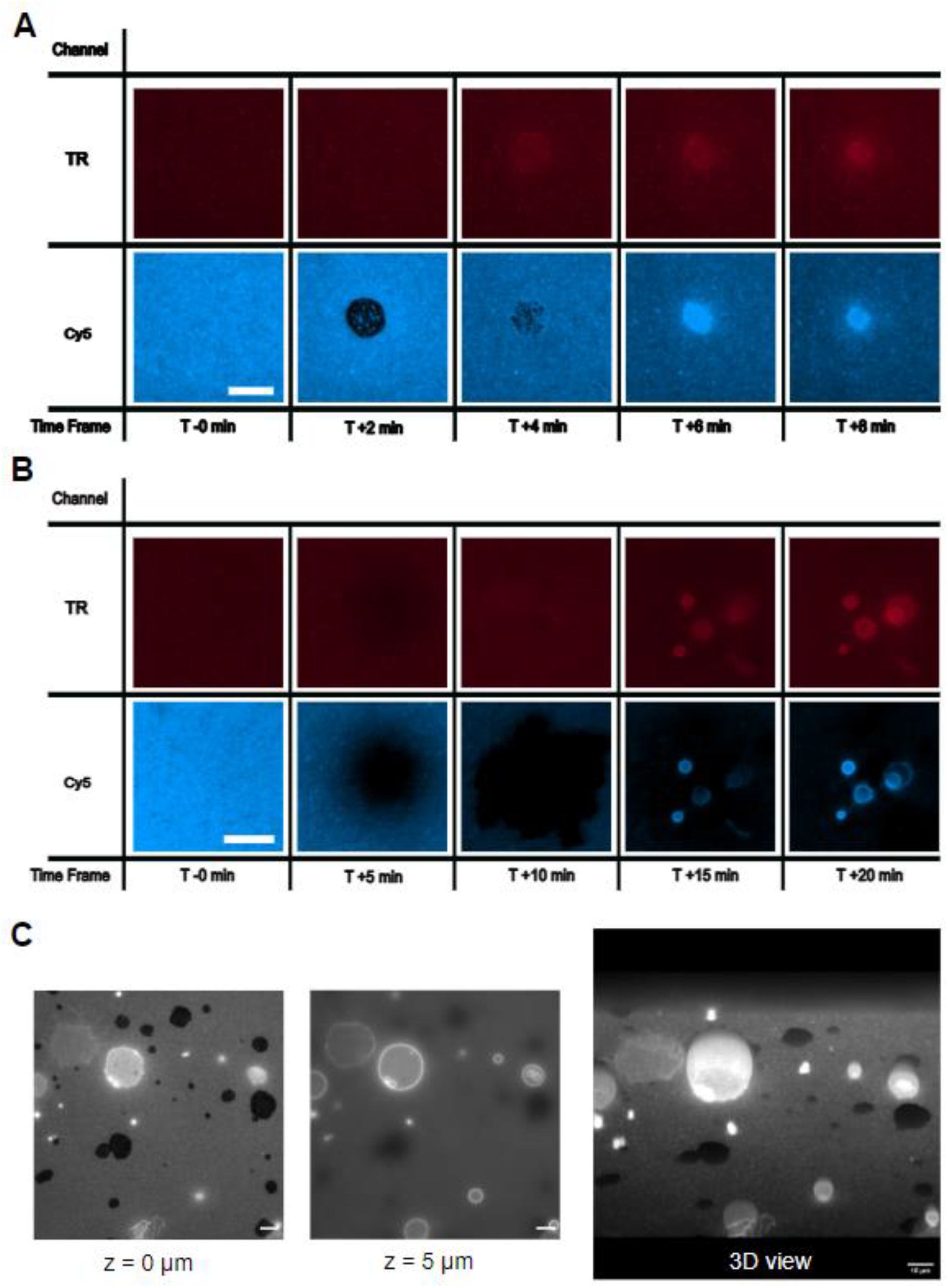
Time lapse movies and 3-dimensional view image of vesicles visible from contact and peptide exchange on the SLB. (A) Colorless GUVs touch and exchange with the SLB and peptide-A647 incubated on the SLB. Interaction is observed right on the surface and time-lapse imges of 0, +2, +4, +6, +8 min are shown. The SLB is viewed in the Texas Red channel distinguished in red. Peptide-A647 is viewed in the Cy5 channel distinguished in blue. (B) Colorless GUVs touch and exchange with the SLB and peptide-A647 incubated on the SLB. Interaction is observed right on the surface and defined vesicles are seen. Time-lapse images of 0, +5, +10, +15 +20 min are shown. The SLB is viewed in the Texas Red channel distinguished in red. Peptide-A647 is viewed in the Cy5 channel distinguished in blue. Scale bars are set to 5 μm. (C) Images taken on the SLB surface and 5 μm above the surface to display the interaction and exchange that took place between GUVs and the SLB. 21 sections of z-stack images for every 1 μm were used to reconstruct the 3D view. Dark contact sites and peptide exchanged vesicles are clearly visible by peptide-A647 fluorescence. Scale bars are 10 μm.

We hypothesize this is by forming hemi-fusion lipid bridges between SLB and GUVs as shown in the schematic of Figure 1A. We saw multiple inhomogeneous fluorescence fluctuations at the contact sites before fluidity was established between GUV and SLB, so we assume majority of the area at the contact sites were involved in hemi-fusion. It is unlikely though each GUV was forming a single hemi-fusion diaphragm spanning the entire area of the entire site with SLB because such single diaphragm formation would involve peptide diffusion only at the very edge of the hemi-fusion site. More likely mechanism is that multiple intermittent hemi-fusion bridges were formed as evidenced by visible diffusion of peptides at the contact site. Our time lapse movies were taken every 30 s, and lipid/peptide exchange typically occurred within a time frame once started, this timescale is close to the natural diffusion rate of the lipids and proteins in SLB [40]. In no case did we observe GUVs collapsing to become a continuous bilayer with the original SLB nor had the SLB budded out to create new vesicles. We did see however, some cases where the landed GUVs fragmented into smaller vesicles before the exchange happened (Figure 3B). We assume this is a large GUV being fragmented due to GUV-SLB interaction followed by smaller GUVs exchanging lipids and peptides by forming hemi-fusion bridges. Many vesicles remained in position suggesting such hemi-fused structure may stay, but there were also some vesicles that were free to diffuse in solution indicating reversing the hemi-fusion state of an individual GUV to leave the contact site is possible resembling the “kiss an run” mechanism, temporary fusion of a vesicle at the contact site followed by fission, in the trafficking of neurotransmitter and endocytosis [41].

Figure 3C clearly summarizes the observation. Contact sites excluded peptides making them clearly visible, but only some of the contacts led to successful exchange of peptides making the GUVs visible by the peptide fluorescence. When such an exchange took place, the outcome was clearly visible. FRAP images showing fluidity of SLB-GUV hemi-fused site can be found in Supplementary S3. GUVs mostly retained the morphologies as vesicles sitting on SLB instead of being entirely fused with SLB. At this point, further major change was seldom observed.

### 3.4. Larger protein cargos do not transfer between SLB and GUV

Hinted by the lipid exchange resulting in spontaneous transfer of small peptides between SLB and GUV, we tested if larger peripheral proteins comparable to the size of a typical protein cargo of tens of kD with addition of a few small protein modifiers on the lipid membranes may be transferred the same way [42]. We performed a similar experiment with SUMO3-GFP cargo instead of small peptides anchored to the SLB. SUMO protein is a small protein modifier that is often attached to a target protein to affect their metabolism and function and GFP is a fluorescent reporter commonly used in cellular imaging [36]. Overall, this cargo is 77 kD (27kD EGFP + 50kD SUMO3 with tags) which is much greater in size compared to the small LAT peptide (4kD). They were anchored to the lipid membrane using an identical chemistry of his-tag Ni binding, and were free to diffuse with lipids in the membranes.

As shown in Figure 4, unlike small peptides, these larger membrane cargos failed to spontaneously transfer between SLB and GUV at least not to the level of clearly quantifiable amount. Visual observation showed no apparent protein bound vesicles emerged as a result of the interaction. We interpret this as a result of the bulkiness of protein size preventing larger proteins to diffuse across the lipid bridge stalks between SLB and GUVs. The geometry of bridges between SLB and GUV are speculated to be challenging for bulky membrane proteins to freely diffuse due to highly curved shape that is necessary to achieve intermittent lipid stalks between proximal bilayers.

**Figure 4.**
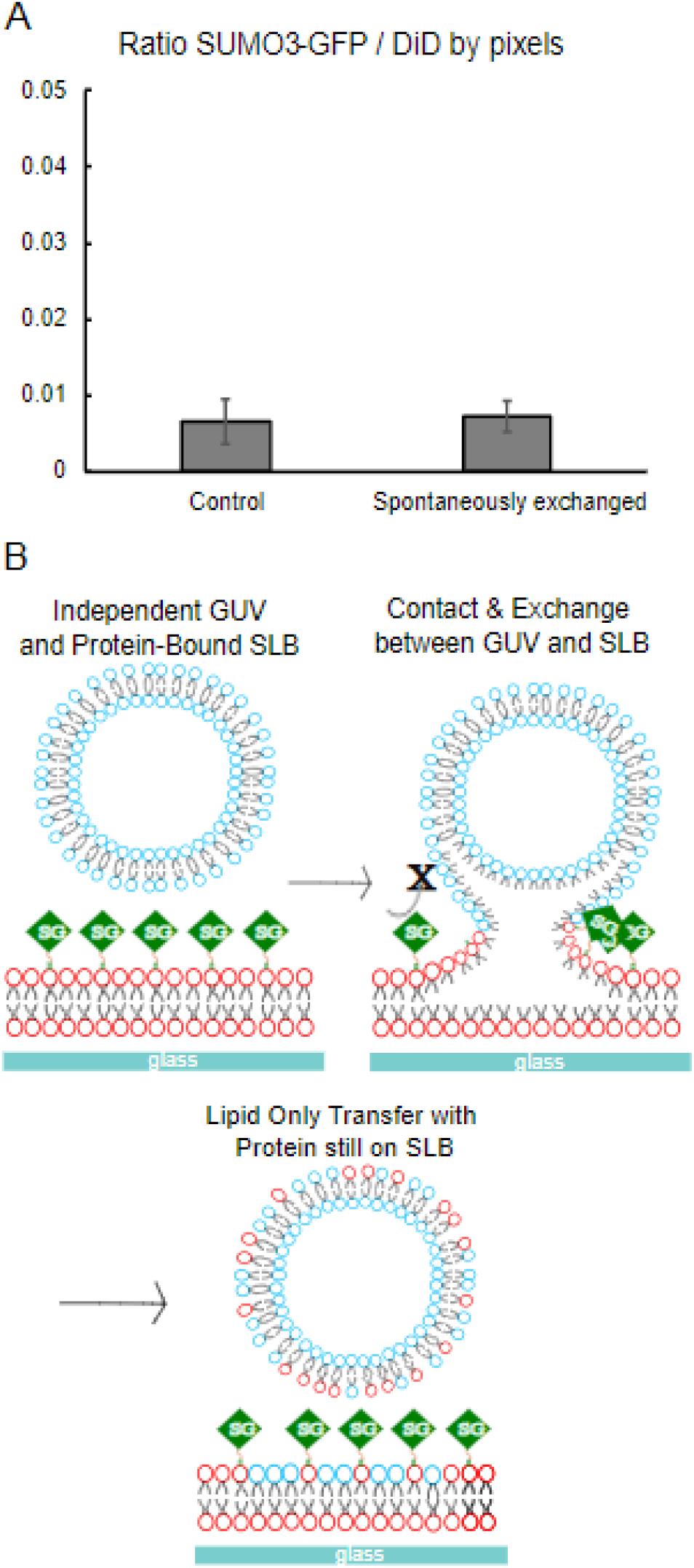
SUMO3-GFP exchange experiments. (A) Statistical distribution by pixels of the ratio of SUMO3-GFP to DiD in a negative control setting compared to spontaneous exchange. There is no significant difference between the control and the spontaneous exchange samples. (B) Schematic of the SLB to GUV exchange with SUMO3-GFP anchored onto the SLB. Bulky SUMO3-GFP proteins (76kD) fail to make transfer from SLB to GUV. The lipid composition for the GUV was 64.9% DOPC, 30% DOPE, 5% DOPS, and 0.1% DiD. The lipid composition for the SLB was 64.9% DOPC, 30% DOPE, 5% Ni-DGS, and 0.1% TR-DHPE.

### 3.5. Facilitating further lipid exchange by cationic shock does not enhance the efficiency of the peptide exchange

To further test the hypothesis of small peptides being transferred by passive diffusion along with lipids, we used a fusogenic condition of highly charged cationic shock by trivalent La3+. By the cationic shock, we catalyzed the lipid exchange between SLB and GUV to monitor if the peptide exchange was enhanced by the increased amount of lipid exchange. It was the choice of fusogenic condition because La3+ is known to induce fusion between lipid bilayers [20], and a previous report showed divalent cation Ca2+ could catalyze lipid exchange between SLB and GUV [38].

As quantified in Figure 5A cationic shock did increase the amount of lipid exchange between SLB and GUVs as evidenced by the increased TR/DiD pixel ratio to 0.14. We could occasionally see induced fusion among GUVs as well at this condition (Supplementary S2). However, against the assumption of catalyzing lipid exchange inducing more peptide exchange, the amount of peptide exchange remained unchanged before and after the cationic shock (Figure 5B). Related to the larger protein exchange experiments, this result suggests that while lipid exchange is a prerequisite for the transfer of membrane-bound peptides, lipid exchange alone is an insufficient condition for peptide exchange. In this case, cationic shock catalyzed the lipid exchange between SLB and GUV, but could not further catalyze the exchange of peripheral peptides.

**Figure 5.**
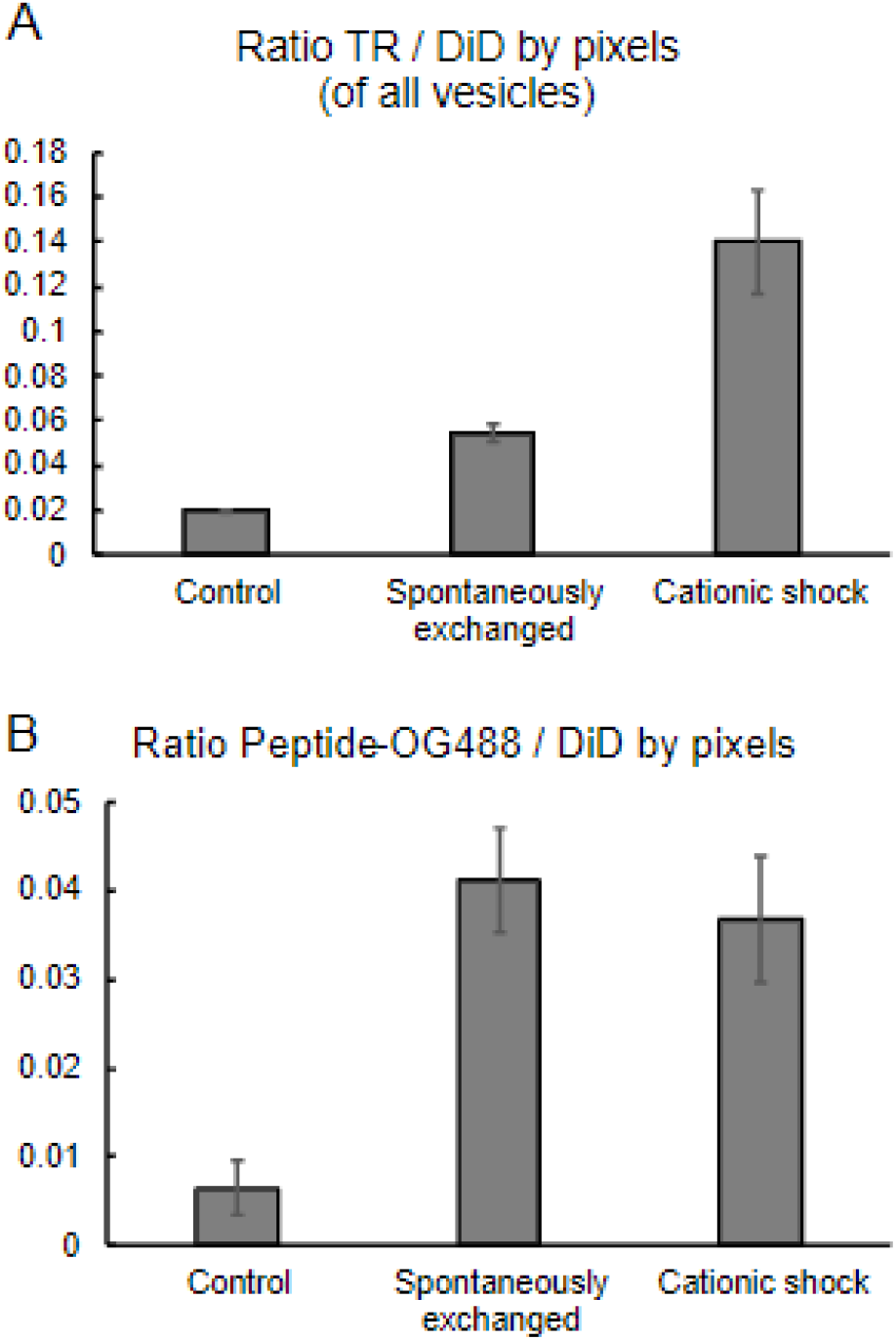
Cationic shock experiments to catalyze lipids exchange. (AB) Statistical distribution by pixels of the ratio of TR-DHPE vesicles to DiD vesicles in a negative control setting compared to spontaneous exchange and cationic shock with lanthanide. Cationic shock with La^3+^ has the greatest effect on the interaction, increasing the proportion of TR-DHPE vesicles to DiD vesicles facilitating the lipid exchange. (A) Statistical distribution by pixels of the ratio of LAT-OG488 to DiD in a negative control setting compared to spontaneous exchange and cationic shock with La^3+^. While cationic shock does catalyze the lipid exchange, it does not result in further exchange of peptides between SLB and GUVs.

### 3.6. Current model of the peptide transfer

Our work reports a case of synthetic lipid membrane system trafficking membrane peptides spontaneously under the neutral pH 7.4 and physiological ionic strength of 150 mM NaCl condition. The work is part of our group’s effort to construct a minimal synthetic membrane system to perform an efficient trafficking of membrane bound proteins. Our result serves an example platform where membrane bound peptide can be transferred between SLB and GUV, and also shed hint on the mechanism of membrane protein transfer between relatively large (>1 μm2) lipid bilayers.

Our hypothetical model for the peptide transfer is that when SLB and GUVs form contact sites, occasional lipid hemi-fusion bridges form. Lipids can thermally diffuse across the bridge, and membrane anchored protein cargos can cross the bridge when their sizes are small enough to diffuse through the highly curved lipid bridges. This is evidenced by the experiment where larger

SUMO3-GFP cargos could not be transferred while smaller LAT peptides could be transferred between the membranes. Catalyzing the lipid exchange by cationic shock could not further enhance the chance of peptide transfer and this may suggest that catatonically catalyzed lipid exchange occurs by a different mechanism that does not provide a chance for small peripheral peptides to transfer.

Our work suggests an experimental platform to establish peptide exchange between SLB and GUVs, and provides a hypothetical model of peptide transfer, but caution should be taken interpreting the results. Firstly, even though the observed spontaneous exchange was reproducible and clearly observable in all our experiments with various combinations of dyes used, not every GUV showed such exchange after contacting with SLBs. We tested a prolonged incubation assuming this was merely due to a very slow kinetic process, but we could not find evidence that longer incubation led to successful peptide transfer of every vesicle. (data now shown) It may suggest that certain vesicles are in the ‘right’ condition to spontaneously react with SLB which is dependent on unknown parameters such as membrane tension, or exact lipid composition. Secondly, Ni-DGS was used as a necessary component to anchor peptides and proteins by his-tag chelation. We speculate Ni-chelation chemistry might be influenced by the cationic shock condition affecting the final outcome of the cationic shock experiments in Figure 5. Thirdly, our model can phenomenologically explain what structural change was undergoing between the two bilayers but does not unambiguously pinpoint the major driving force allowing this exchange to occur between GUV and SLB at the condition tested.

## 4. Conclusion

We report a case of synthetic lipid membrane platform capable of trafficking small membrane bound peptides from SLB to GUVs. A series of experiments support the model that the transfer occurs by formation of hemi-fused lipid stalks between GUV and SLB that are highly curved limiting diffusion of larger protein cargos. It suggests peripheral peptides may transfer between the membranes by occasional hemi-fusion interaction and the transfer is limited by the geometry of the lipid bridge which is hypothesized to be highly curved. Our model provides an insight for a simple mechanism of lipid bilayers to interact at a neutral saline buffer condition to transfer membrane proteins spontaneously and serves an example of synthetic system capable of transferring small peptides between two lipid bilayers with a relatively large contact area (> 1 μm2). Potential future work includes exploring the conditions of buffers and lipid compositions affecting the exchange with the goal of pinpointing the major driving force causing the hemi-fusion, and single molecule tracking study to monitor the movement of peptide molecules at a higher spatial and temporal resolution to monitor the movement of peptide molecules diffusing across the hemi-fused stalks. Eventually, discovering conditions where larger protein cargos can be transferred is expected to provide us a useful functional platform of vesicular trafficking that is compatible with dynamic high-resolution imaging by optical microscopy.

## Abbreviations

A647: Alexa fluor 647
CLSM: Confocal laser scanning microscope
DiD: 1,1′-dioctadecyl-3,3,3′,3′-tetramethylindodicarbocyanine
DOPC: 1,2-Dioleoyl-sn-glycero-3-phosphocholine
DOPE: 1,2-dioleoyl-sn-glycero-3-phosphoethanolamine
DOPS: 1,2-dioleoyl-sn-glycero-3-phospho-L-serine
FRAP: Fluorescence bleaching after photobleaching
GFP: Green fluorescence protein
GUV: Giant unilamellar vesicle
LAT: Linker for activation of T-cell
Ni-DGS: Nickel bound 1,2-Dioleoyl-sn-glycero-3-[(N-(5-amino-1-carboxypentyl)iminodiacetic acid)succinyl]
OG488: Oregon green 488
SLB: Supported lipid bilayer
SUMO: Small Ubiquitin-like modifier
TR-DHPE: Texas Red-1,2-Dihexadecanoyl-sn-Glycero-3-Phosphoethanolamine

## Acknowledgements

The authors would like to thank Dr. Laying Wu of the Microscopy and microanalysis research laboratory at Montclair state university for the microscope training and facility maintenance.

## Appendix A. Supplementary data

Supplementary data to this article can be found online.

