## Supplementary materials for "Spontaneous transfer of small peripheral peptides between supported lipid bilayer and giant unilamellar vesicles"

### Supporting Information for the article: "Spontaneous transfer of small peripheral peptides between supported lipid bilayer and giant unilamellar vesicles"

Emanuela Efodili[1], Ashlynn Knight[2], Maryem Mirza[3], Cedric Briones[1], and Il-Hyung Lee[1*]

[1]Department of Chemistry and Biochemistry, Montclair State University, Montclair, NJ 07043, USA, [2]Department of Biology, Montclair State University, Montclair, NJ 07043, USA, [3]College of humanities and social sciences, Montclair State University, Montclair, NJ 07043.

**The file includes:**

S1-S2. Supplementary figures

S3. Raw code of the image analysis

#### S1. Supplementary figure 1

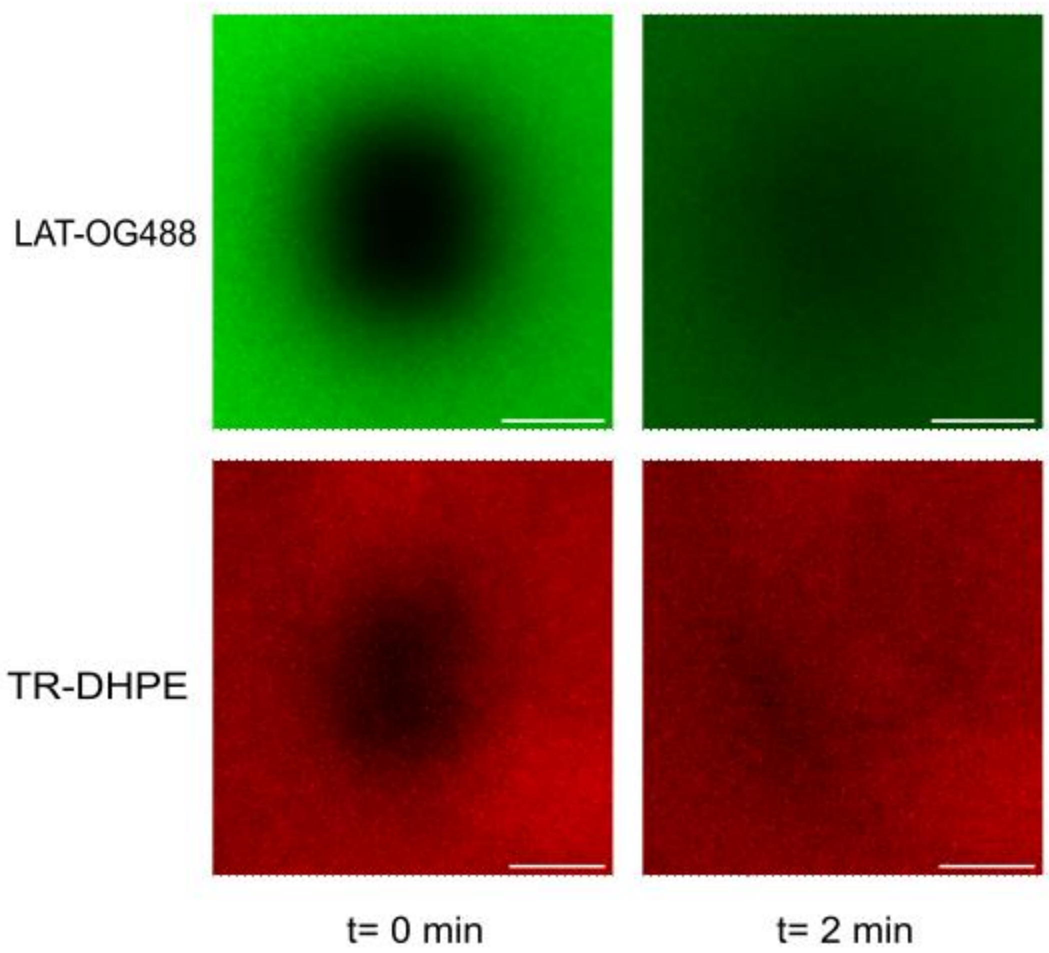


**Supplementary figure 1**. Fluorescence recovery after photobleaching (FRAP) to verify lipid bilayer fluidity. The peptide LAT-OG488 forms a layer on top of the SLB by anchoring onto Ni-DGS present in the supported bilayer. LAT-OG488 is represented in the green GFP channel, where the membrane appears mobile and fluidic, evidenced by disappearance of bleached area by full fluorescence recovery. SLB composition was 64.9% DOPC, 30% DOPE, 5% Ni-DGS, and 0.1% TR-DHPE. The SLB is represented in the red Texas Red channel, where the membrane appears mobile and fluidic, and full recovery is observed. Scale bars are set to 10 μm. Images acquired by wide field fluorescence microscopy.

#### S2. Supplementary figure 2

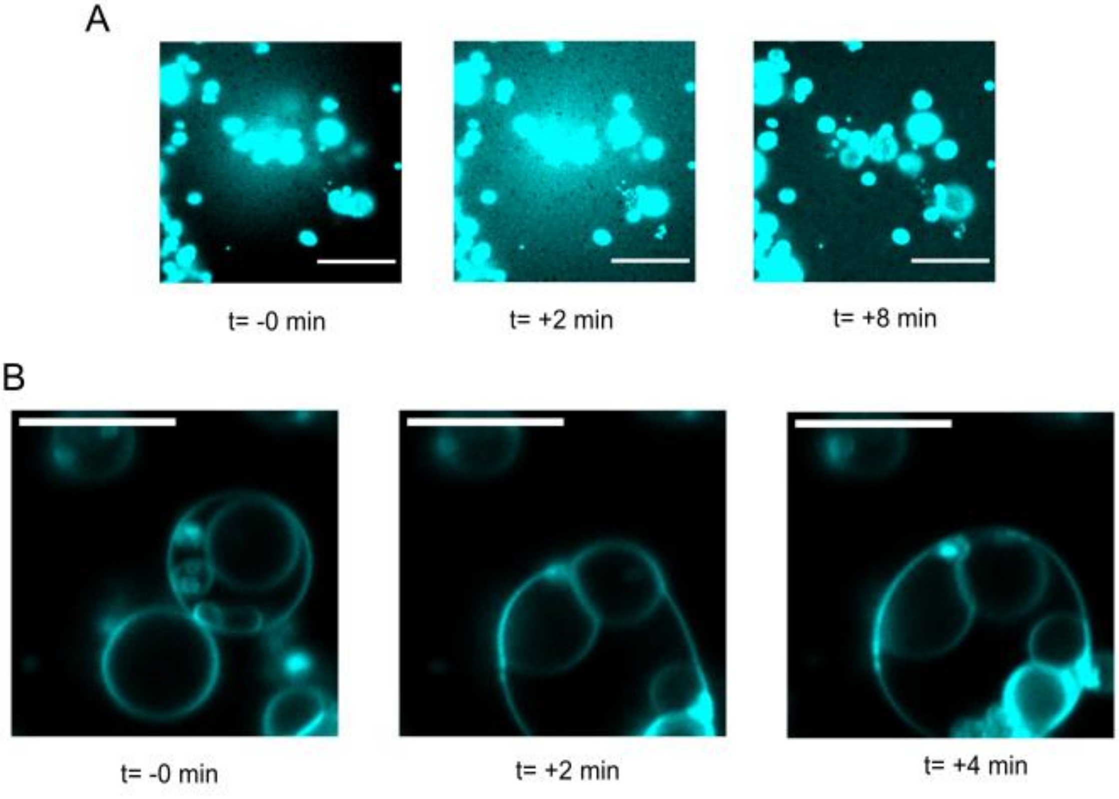


**Supplementary figure 2**. Example images showing the effect of cationic shock. (A) Cationic shock catalayzed the lipid exchange. TR-DHPE fluorescence, originally in GUVs diffuses across the SLB as a visible wave of fluorescence intensity propagating outward. (B) Cationic shock of the sample induced GUV to GUV fusion. Two independent vesicles were fused into one. 1 mM $La^{3+}$ was gently introduced to the system. The lipid composition of the GUVs was 94.9% DOPC, 5% DOPS, and 0.1% TR-DHPE. The lipid composition of the SLB was 94.9% DOPC, 5% Ni-DGS, and 0.1% DiD. Scale bar set to 10 μm.

#### S3. Supplementary figure 3

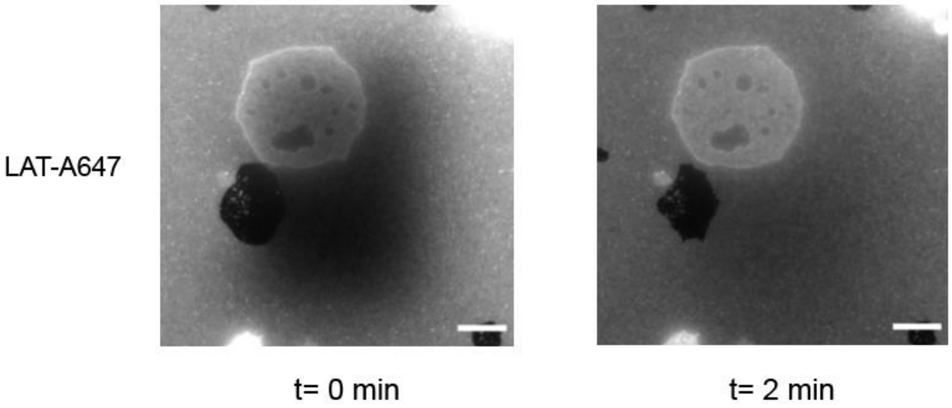


**Supplementary figure 3**. Fluorescence recovery after photobleaching (FRAP) to verify lipid bilayer fluidity including the hemi-fusion site. Both original SLB and the GUV bottom membrane in contact are fully recovering the fluorescence suggesting that the hemi-fused bridge was established between GUV and SLB. SLB composition was 65% DOPC, 30% DOPE, 5% Ni-DGS, and GUV composition was 65% DOPC, 30% DOPE, 5% DOPS. LAT-A647 fluorescence signal is from peptides anchored to the SLB by Ni-His tag binding. Experimental condition of the incubation was as described in the main text. Scale bars are set to 10 μm. Images acquired by wide field fluorescence microscopy.

#### S4. Raw code of the image analysis

This code can be saved as any name (e.g. MaskInt2.m) to run by GNU Octave. Ver.6.4.0 was used for the analysis and was confirmed to run this code with no errors. Images to analyze should be under the folder named "Sample". "Sample\A" should contain the reference images of Channel A, "Sample\B" and "Sample\C" should contain matching images for Channel B and C each. Matching images should be saved as .tif files with same names as the image files will be recognized by alphabetical order. Analysis output will be saved in the folder "Analysis". Comments were color-coded for readability.

The raw code starts below:

```
%%%%%%%%%%%%%%%%%%%%%%%%%%%%%%%%%%%%%%%%%%%%%%%%%%%%%%%%%%%%%%%%%%
% GUV binding intensity Analysis program
%%%%%%%%%%%%%%%%%%%%%%%%%%%%%%%%%%%%%%%%%%%%%%%%%%%%%%%%%%%%%%%%%%
% Important parameters to set by users

pkg load image;

StckN = 3; %%stack number to analyze (from the bottom)

autoBackground = 4; %automatic estimation of background intensity by most probable
                    %vlaue 1 to use mode estimation, value 2 for median estimation
                        %3 for triangle thresholding, requires triangle_th.m by Bernard Panneton
                    %4 for a defined manual calculation

backgroundIntA = 250;  %manual input of the backgroudn intensity in case not using auto estimation
backgroundIntB = 50;
backgroundIntC = 50;

cutNoise = 2; % pixels with intensities below backgroundInt x cutNoise will not be masked

ignoreSat = 1; %1 - to ignore the saturated pixels
satIntensity = 4095; %saturation intensity

saveCircles = 1; %to save mask information

%%%%%%%%%%%%%%%%%%%%%%%%%%%%%%%%%%%%%%%%%%%%%%%%%%%%%%%%%%%%%%%%%%
%%%%%%%%%%%%%%%%%%%%%%%%%%
%PathName=uigetdir('Choose file folers containing directory (two colors)');
PathName = './Sample/';
filelistA = dir([PathName, '/A/*.tif']); % reference channel
filelistB = dir([PathName, '/B/*.tif']); % binding or changing channel
filelistC = dir([PathName, '/C/*.tif']);
if length(filelistA) != length(filelistB)
  disp('number of stacks mismatch in two channels');
  exit;
endif
PathOut = [PathName '\Analysis'];
if exist([PathName '\Analysis']) != 7
  mkdir(PathOut);
endif
%%%%%%%%%%%%%%%%%%%%%%%%%%%%%%%%%%%%%%%%%%%%%%%%%%%%%%%%%%%%%%%%%%
%%%%%%%%%%%%%%%%%%%%%%%%
intensity = zeros(length(filelistA),21);
  %Intensity analysis
  #[total_intensityA     number_of_pixelsA       average_intensityA
  # average_backgroundA   int-backgroundA
  #total_intensityB     number_of_pixelsB       average_intensityB
```

```
  # average_backgroundB   int-backgroundB
  #total_intensityC     number_of_pixelsC       average_intensityC
  # average_backgroundC   int-backgroundC
  # normbalizedB/A    normbalizedC/A       normbalizedC/B
  # maskedPixelsB/A    maskedPixelsC/A       maskedPixelsC/B]

for index=1:1:length(filelistA)

  fnameA = [PathName '\A\' filelistA(index).name];
  fnameB = [PathName '\B\' filelistB(index).name];
  fnameC = [PathName '\C\' filelistC(index).name];
  imgAinfo = imfinfo(fnameA);
  record = []; %variable to store each image data
  GUV = []; %variable to track each GUV
  disp(['File ', [num2str(index)],' open...']);

  for indexStack=StckN:1:StckN %just one stack image analysis

  imgA = imread(fnameA, indexStack);
  imgB = imread(fnameB, indexStack);
  imgC = imread(fnameC, indexStack);
  imgA = uint16(imgA); %casting added to avoid data type confusion in thresholding
  imgB = uint16(imgB);
  imgC = uint16(imgC);

  %%%%%%%%%%

  %background noise level analysis
  if autoBackground == 1
    backgroundIntA = mode(imgA(:));
    backgroundIntB = mode(imgB(:));
    backgroundIntC = mode(imgC(:));
  elseif autoBackground == 2
    backgroundIntA = median(imgA(:));
    backgroundIntB = median(imgB(:));
    backgroundIntC = median(imgC(:));
  elseif autoBackground == 3
    [lehisto x] = imhist(imgA,11107); %triangle thresholding
    backgroundIntA = 65535 * triangle_th(lehisto, 11107);
    [lehisto x] = imhist(imgB,11107); %triangle thresholding
    backgroundIntB = 65535 * triangle_th(lehisto, 11107);
    [lehisto x] = imhist(imgC,11107); %triangle thresholding
    backgroundIntC = 65535 * triangle_th(lehisto, 11107);
  elseif autoBackground == 4
    backgroundIntA = median(imgA(:))+ max(imgA(:))/10;
    backgroundIntB = median(imgB(:))+ max(imgB(:))/10;
    backgroundIntC = median(imgC(:))+ max(imgC(:))/10;
  endif

  ####create mask image for output###
  imgAout = imgA;
  imgAout(:,:) = 0;
  imgBout = imgB;
  imgBout(:,:) = 0;
  imgCout = imgC;
  imgCout(:,:) = 0;
  #############################



  #################
```

```
    for ix=1:1:size(imgA,2)
      for iy=1:1:size(imgA,1)
        if int32(imgA(iy, ix)) <= backgroundIntA * cutNoise
          continue;
        elseif (ignoreSat == 1) && (int32(imgA(iy, ix)) >= satIntensity)
          continue;
        endif
          intensity(index,1) += int32(imgA(iy, ix));
          intensity(index,2) += 1;
          imgAout(iy, ix) = 255;
          if int32(imgB(iy, ix)) <= backgroundIntB * cutNoise
            continue;
          else
            intensity(index,6) += int32(imgB(iy, ix));
            intensity(index,7) += 1;
            imgBout(iy, ix) = 255;
          endif
          if int32(imgC(iy, ix)) <= backgroundIntC * cutNoise
            continue;
          else
            intensity(index,11) += int32(imgC(iy, ix));
            intensity(index,12) += 1;
            imgCout(iy, ix) = 255;
          endif
      endfor
    endfor
    intensity(index,3) = intensity(index,1) / intensity(index,2);
    intensity(index,4) = backgroundIntA;
    intensity(index,5) = intensity(index,3) - backgroundIntA;
    intensity(index,8) = intensity(index,6) / intensity(index,7);
    intensity(index,9) = backgroundIntB;
    intensity(index,10) = intensity(index,8) - backgroundIntB;
    intensity(index,13) = intensity(index,11) / intensity(index,12);
    intensity(index,14) = backgroundIntC;
    intensity(index,15) = intensity(index,13) - backgroundIntC;

    intensity(index,16) = intensity(index,10) / intensity(index,5);
    intensity(index,17) = intensity(index,15) / intensity(index,5);
    intensity(index,18) = intensity(index,15) / intensity(index,10);

    intensity(index,19) = intensity(index,7) / intensity(index,2);
    intensity(index,20) = intensity(index,12) / intensity(index,2);
    intensity(index,21) = intensity(index,12) / intensity(index,7);

  disp(['Image', num2str(index), ' IntensityA:', num2str(intensity(index,5)),' IntensityB:', num2str(intensity(index,10)), '
IntensityC:', num2str(intensity(index,15))]);
  disp(['Image', num2str(index), ' PixelsA:', num2str(intensity(index,2)),' PixelsB:', num2str(intensity(index,7)), '
PixelsC:', num2str(intensity(index,12))]);


  endfor % stack loop


  %%%%%Save relevant information%%%%
  %%header and clean output
  OutName = strrep(filelistA(1).name,'.tif','.txt');
  %dlmwrite([PathOut '\' OutName],toSave,'delimiter','\t')
    fid = fopen([PathOut '\' OutName], 'wt');
    fprintf(fid, '%s\t%s\t%s\t%s\t%s\t%s\t%s\t%s\t%s\t%s\t%s\t%s\t%s\t%s\t%s\t%s\t%s\t%s\t%s\t%s\t%s\n',
'total_intensityA', 'num_of_pixelsA', 'avg_intA', 'background_intA', 'adjusted_intA', 'total_intensityB', 'num_of_pixelsB',
```

```
'avg_intB', 'background_intB', 'adjusted_intB', 'total_intensityC', 'num_of_pixelsC', 'avg_intC', 'background_intC',
'adjusted_intC', 'intB/intA', 'intC/intA', 'intC/intB', 'pixB/pixA', 'pixC/pixA', 'pixC/pixB');  % header
    fclose(fid);
    dlmwrite([PathOut '\' OutName],intensity,'delimiter','\t','-append');

  if saveCircles == 1
        circleFileName = ['A_' filelistA(index).name];
        %H = getframe(figure(indexStack));
        imwrite(imgAout, [PathOut '/' circleFileName]);
        circleFileName = ['B_' filelistA(index).name];
        imwrite(imgBout, [PathOut '/' circleFileName]);
        circleFileName = ['C_' filelistA(index).name];
        imwrite(imgCout, [PathOut '/' circleFileName]);
  endif
  %%%%%%%%%%%%%%%%%%%%%%%%%%%%%%%%%%
  %%%%%%%%%%%%%%%%%%%%%%%%%%%%%%%%%%%%%%%%%%%%%%%%%%
endfor
```
